# Quantifying innervation facilitated by deep learning in wound healing

**DOI:** 10.1101/2023.06.14.544960

**Authors:** Abijeet Singh Mehta, Sam Teymoori, Cynthia Recendez, Daniel Fregoso, Anthony Gallegos, Hsin-Ya Yang, Roslyn Rivkah Isseroff, Min Zhao, Marcella Gomez

## Abstract

The peripheral nerves (PNs) innervate the dermis and epidermis, which have been suggested to play an important role in wound healing. Several methods to quantify skin innervation during wound healing have been reported. Those usually require multiple observers, are complex and labor-intensive, and noise/background associated with the Immunohistochemistry (IHC) images could cause quantification errors/user bias. In this study, we employed the state-of-the-art deep neural network, DnCNN, to perform pre-processing and effectively reduce the noise in the IHC images. Additionally, we utilized an automated image analysis tool, assisted by Matlab, to accurately determine the extent of skin innervation during various stages of wound healing. The 8mm wound is generated using a circular biopsy punch in the wild-type mouse. Skin samples were collected on days 3,7,10 and 15, and sections from paraffin-embedded tissues were stained against pan-neuronal marker- protein-gene-product 9.5 (PGP 9.5) antibody. On day 3 and day 7, negligible nerve fibers were present throughout the wound with few only on the lateral boundaries of the wound. On day 10, a slight increase in nerve fiber density appeared, which significantly increased on day 15. Importantly we found a positive correlation (R-^2^ = 0.933) between nerve fiber density and re-epithelization, suggesting an association between re-innervation and re-epithelization. These results established a quantitative time course of re-innervation in wound healing, and the automated image analysis method offers a novel and useful tool to facilitate the quantification of innervation in the skin and other tissues.

## 1. Introduction

Wound regeneration is a complex process that is regulated by orchestrated mechanisms, influenced by chemical, cellular, and molecular factors^1–2^. The healing process begins at the time of injury and eventual maturation could continue for months or even years until the wound completely heals and is structurally and functionally similar to uninjured skin^3^. The four overlapping phases of wound healing are homeostatic, inflammatory, proliferative, and remodeling (Figure 1). The homeostatic phase lasts a few hours producing a fibrin plug followed by an inflammatory phase, which can last between hours and days, during which aggregated platelets and cells release pro-inflammatory mediators^4^. The early inflammatory phase is succeeded by the proliferative phase lasting a few weeks during which macrophages and fibroblast cells invade the wound bed forming granulation tissue and active migration of the wound epithelial cells occurs^5^. The last phase of wound healing is the remodeling phase, which is characterized by proliferative cell apoptosis, adjustment of extracellular matrix (ECM), and replacement of type 3 with type 1 collagen in the dermis. It can last for weeks to years^3^.

**Figure 1:**
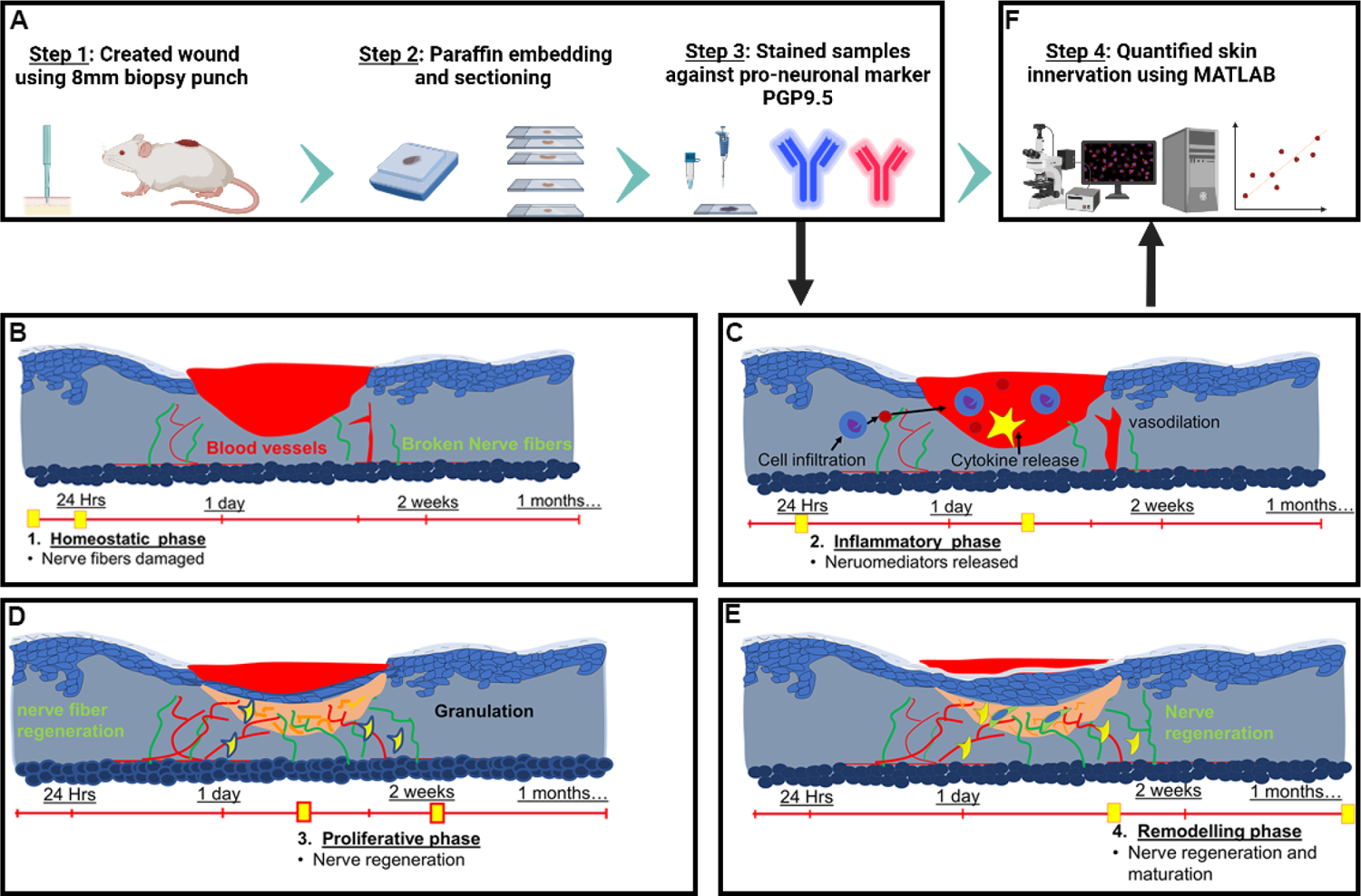
The experimental design and schematic depicting methodology to quantify skin innervation. (A) A biopsy punch of 8mm in diameter is used to create the wound, and skin samples are collected and fixed on days 3, 7,10 and 15. After fixation, the wounded tissue is paraffin-embedded and sectioned (5 μm thickness) for immunofluorescence analysis against PGP9.5 protein, a neuron-specific marker. (B-E) Illustration portraying different stages of wound healing (B) Homeostatic phase last a few hours during which nerve fibers in the wound bed are damaged followed by (C) Inflammatory phase that can last between hours and days. (D) The proliferative phase lasts a few weeks during which re-innervation might be initiated and (E) during the remodeling phase wound matures and can last between weeks to years. In our study, we choose to quantify skin innervation at days 3, 7, 10 and 15 as an attempt to cover all phases of wound healing. (F) The immunohistochemistry (IHC) samples are analyzed using automated Matlab-assisted tools aided by DnCNN-based image denoising.

During healing phases, a strong interaction between the nervous system and skin involving a variety of neuromodulators, cytokines, hormones, and other effector molecules has been reported^6–7^. The nervous system can be influenced both at the local and central levels by the stimuli at the skin and vice versa. The brain can alter skin function during the pathophysiological state and skin can modulate the nervous system by releasing a variety of neuropeptides^8^. The cutaneous nerves therefore positively affect all the stages of wound healing^9^. Numerous neuropeptides e.g. substance P (SP) released from cutaneous nerves have been reported to activate vital mechanisms during the inflammatory phase^10^. Similarly, neuropeptides released by cutaneous nerves influence the proliferation phase. They can promote the proliferation of fibroblasts, keratinocytes, and endothelial cells by stimulating DNA synthesis, can stimulate angiogenesis, support granulation tissue remodeling, and many more ^11–15^. The effect of innervation on the remodeling stage has also been studied in past. It has been demonstrated that a significantly higher number of nerve fibers correlate with normotrophic scars in comparison to hypertrophic scars during the remodeling phase^16–,17^. Hence, the literature strongly suggests a regulatory role for skin nerves in wound healing and any impairment in skin innervation is one of the leading causes of occurrence of chronic wounds e.g., diabetic neuropathy with their foot ulcers and plegias with their sacral and trochanteric pressure sores^18–19^.

Previously, numerous studies have been conducted on quantifying skin innervation^20–26^. However, most such studies are not fully automated, have manual counts of IHC-stained structures that are prone to user errors and variation, require multiple observers, and are complex and labor-intensive. One such example is dendrite analysis involving manually tracing neurons using the simple neurite tracer plug-in of ImageJ software ^21–22^. While this semi-manual approach has been proven effective, it involves identifying the beginning and end points of dendrites and digitally drawing individual branch segments throughout the entire neuron, making it a labor-intensive and time-consuming process. Another example is semi-automated Sholl analysis for quantifying changes in the growth and differentiation of neurons and glia^23^. The method offers several advantages over conventional manual quantification, including faster analysis time and increased statistical sensitivity, however, has some limitations, such as reliance on manual input from the user, which introduces a risk of user error and variation impacting the accuracy and reliability of the results. Additionally, the semi-automated Sholl method is complex and time-consuming to set up initially, which could act as a barrier for researchers who do not have the technical expertise or resources to implement the method effectively. In an effort to quickly, objectively, and reproducibly quantify cutaneous innervation we developed a fully automated Matlab-assisted image analysis tool aided by powerful deep neural network, DnCNN, for pre-processing (de-noising) of the IHC-images. This network can detect and remove high-frequency image artifacts and increase image resolution, noise is minimized, resulting in higher quality images that can be more accurately analyzed. The DnCNN is particularly developed for image processing^27^, and has shown effectiveness in a wide range of applications, including medical imaging^28^.

Utilizing automated Matlab-assisted tool aided with DnCNN we quantified skin innervation during wound healing stages at days 3,7,10 and 15. The data show a positive correlation between the increase in nerve fiber density and re-epithelization.

## 2. Materials and Methods

### 2.1. Animals

All procedures and protocols were reviewed and approved by the institution’s IACUC, and performed at the UC Davis Teaching and Research Animal Care Services (TRACS) facility. Female C57BL/6 (14–15-week-old, 20–24 g) mice were obtained from Jackson Laboratory. The animals were acclimated for one week after transfer from the vendor to the UC Davis vivarium. All female mice used in this experiment were kept in the containment unit of the animal facility, housed in cages with free access to food and water. All experiments were performed in a biosafety cabinet and were done in triplicate (n=3).

### 2.2. Wounding

The animals were sedated with 3–5% isoflurane two days prior to wounding surgery and the dorsal surface was shaved. To remove the remaining hair depilatory cream was applied and removed within 5–10 s of application. For surgery, all mice were anesthetized with isoflurane. <u>Buprenorphine</u> (0.05 mg/kg) was administered subcutaneously prior to the wounding procedure. Iodine and ethanol wipes were used to sterilely prep experimental mice. An 8 mm sterile skin biopsy punch instrument was used to create full-thickness wounds on the dorsal skin as described previously^7, 29^. At the end of the experiment, mice were euthanized by cervical dislocation under 5% isoflurane, and skin samples were collected on days 3,7,10, and 15 post-wounding.

### 2.3. Immunohistochemical analysis of wound tissues

After fixation, the wound tissue was paraffin-embedded and sectioned (5 μm thickness) for immunohistochemical analysis as described previously^30–31^. Primary antibodies against PGP9.5 (Invitrogen, Catalog # PA5-29012), and β-III tubulin (Invitrogen, Catalog # MA5-16308) were used. Donkey anti-Rabbit IgG (H+L) highly Cross-Adsorbed Secondary Antibody, Alexa Fluor™ 594 (Invitrogen, Catalog # A21207), and donkey anti-Mouse IgG (H+L) Highly Cross-Adsorbed Secondary Antibody, Alexa Fluor™ 488 (Invitrogen, Catalog # A21202), respectively were used. VECTASHIELD® Antifade Mounting Media with DAPI (Vector Laboratories, Catalog # H-1200-10) was used to stain nuclei. Slides were imaged with an Olympus FV3000 Confocal Laser Scanning Microscope (Shinjuku City, Tokyo, Japan) as described previously^32–34^ and analyzed using Matlab 2021Image processing.

### 2.4. Quantification

In order to accurately quantify wound healing, several steps are taken to ensure precision and accuracy in counting PGP9.5 positively stained pixels (Figure 2). One of the most important steps is preprocessing the images to remove any unwanted noise or artifacts that could potentially interfere with the analysis. Many factors can contribute to image noise, including non-specific staining, autofluorescence, equipment malfunctions, motion blur, and environmental factors. To eliminate noise from the images, a powerful deep neural network known as the DnCNN network is employed. It is a convolutional neural network (CNN) specifically designed for image denoising^27^. It works by learning to map between noisy images and clean images and then use this mapping to remove noise from new images. The DnCNN network is trained on a large dataset of noisy and clean images and has shown effectiveness in a wide range of applications, including medical imaging^28^. Once the images have been denoised, the next step is to count the PGP9.5 positive pixels in the images. Positive pixels representing innervation were identified by their immunoreactivity to PGP9.5 antibody (red color) (Figures 3 and 4). However, identifying those pixels can be challenging because of the variation in the intensity of fluorescence due to different wound depths, healing stages, expression levels and sizes of expression areas. Additionally, the background intensity can pollute the pixel value counted as nerve fibers and nerve terminals. To overcome these challenges, statistical analysis is used to identify the threshold for immunofluorescence and background value. For the background value, we find the low spectrum values of pixels and remove outliers. After finding the background value, we subtract this value from all pixels, making the pixels in different images more comparable. Next, to find the fluorescence threshold, we analyze the immunoreactivity intensity (red color intensity) of all pixels (subtracted version) in the image and identify the range. Finally, the identified threshold is used to count the number of PGP9.5 positive pixels in the image. Each pixel’s (red fluorescence value) is compared to the threshold, and any pixel with a value above the threshold is considered a positive count for a neuron. This process is repeated for all the pixels in the image, determined together as the total number of neurons. Next to identify nerve fiber density we divided the total number of positive pixels by the area of the wound bed (Figure 3) or the area of the outer edges/ wound center (Figure 4). In addition to measuring nerve fiber density in the whole wound, we also analyzed the density of nerve fibers (positive pixels) separately for both the epidermis and dermis to get a more spatial understanding.

**Figure 2:**
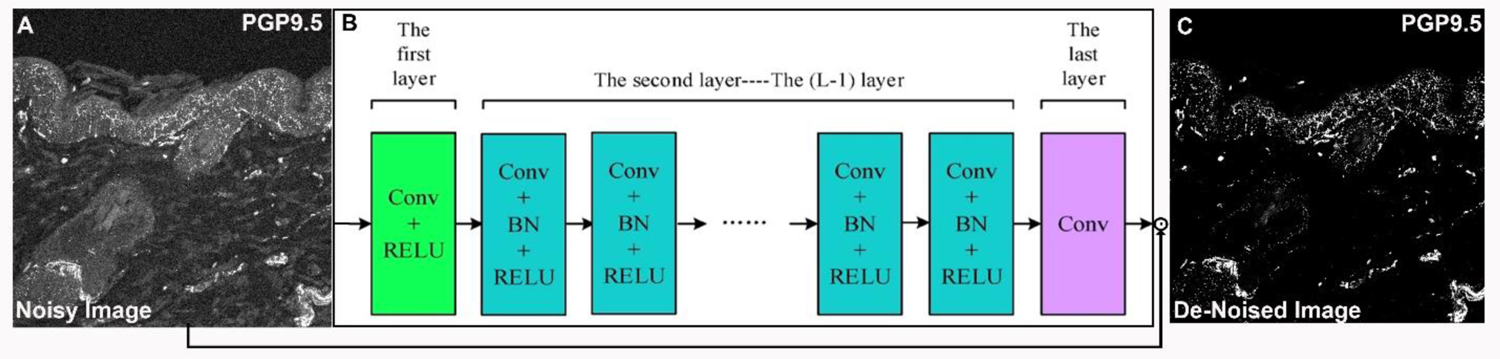
DnCNN network architecture for image denoising. (A) Noisy image as DnCNN input. (B) The DnCNN network architecture consists of multiple convolutional layers. Each convolutional layer includes batch normalization (BN), convolution (Conv), and rectified linear unit (ReLU) layers. The first layer takes the noisy image as input, and the subsequent layers process the image to remove noise. (C) Output image after de-noising.

**Figure 3:**
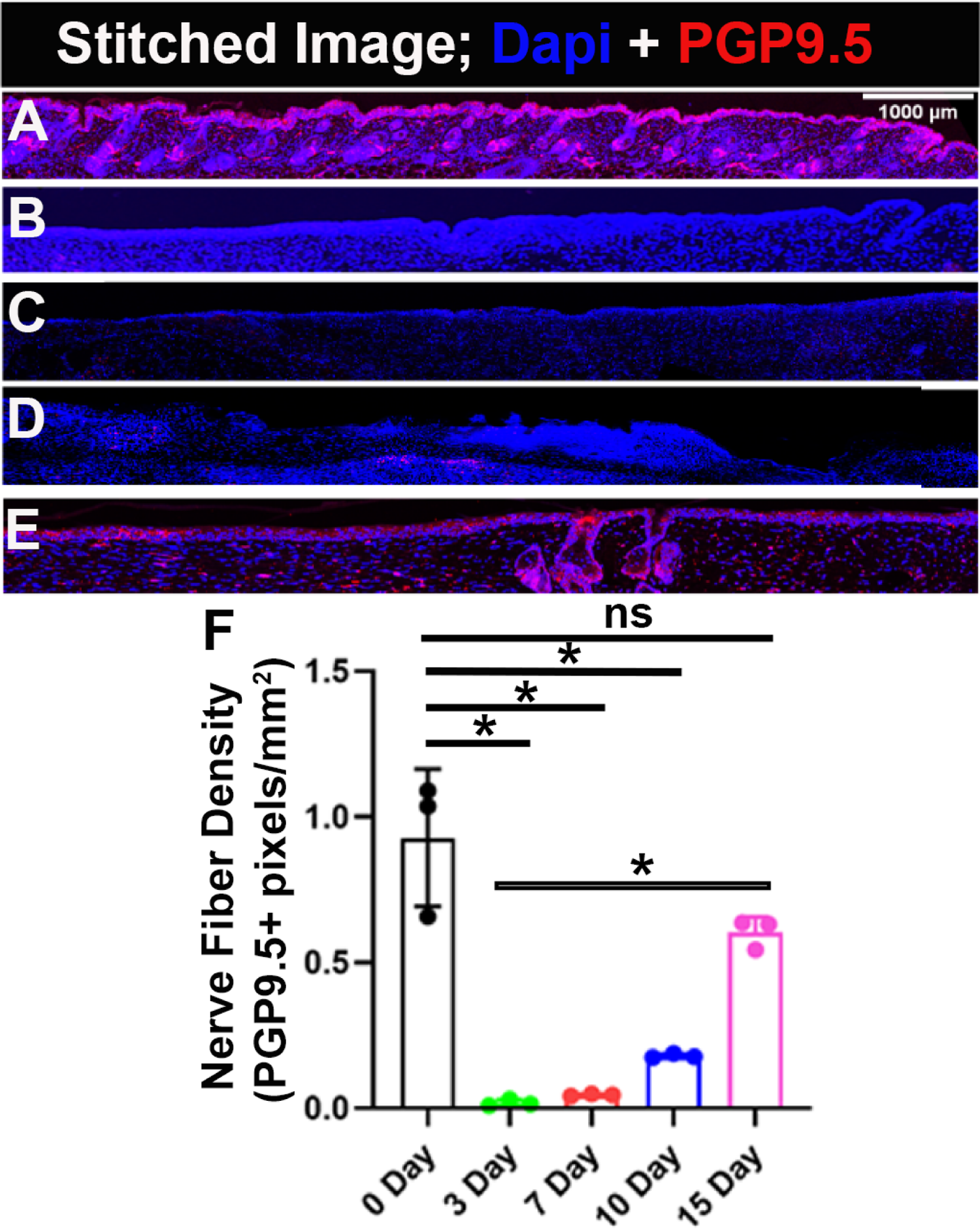
Gradual increase in Re-innervation in the wound bed. PGP9.5 is a pan-neuronal marker and DAPI stains nuclei. (A) Uninjured skin. Skin sample collected on (B) day 3 (C) day 7 (D) day 10 (E) day 15 (F) Quantification of skin innervation for the whole wound bed represented as mean ± SD, n = 3 wounds from three mice in each group, *P < .05, ns is non-significant. The wound bed is recognized by the absence of hair follicles. Scale bar = 1000 μm.

**Figure 4:**
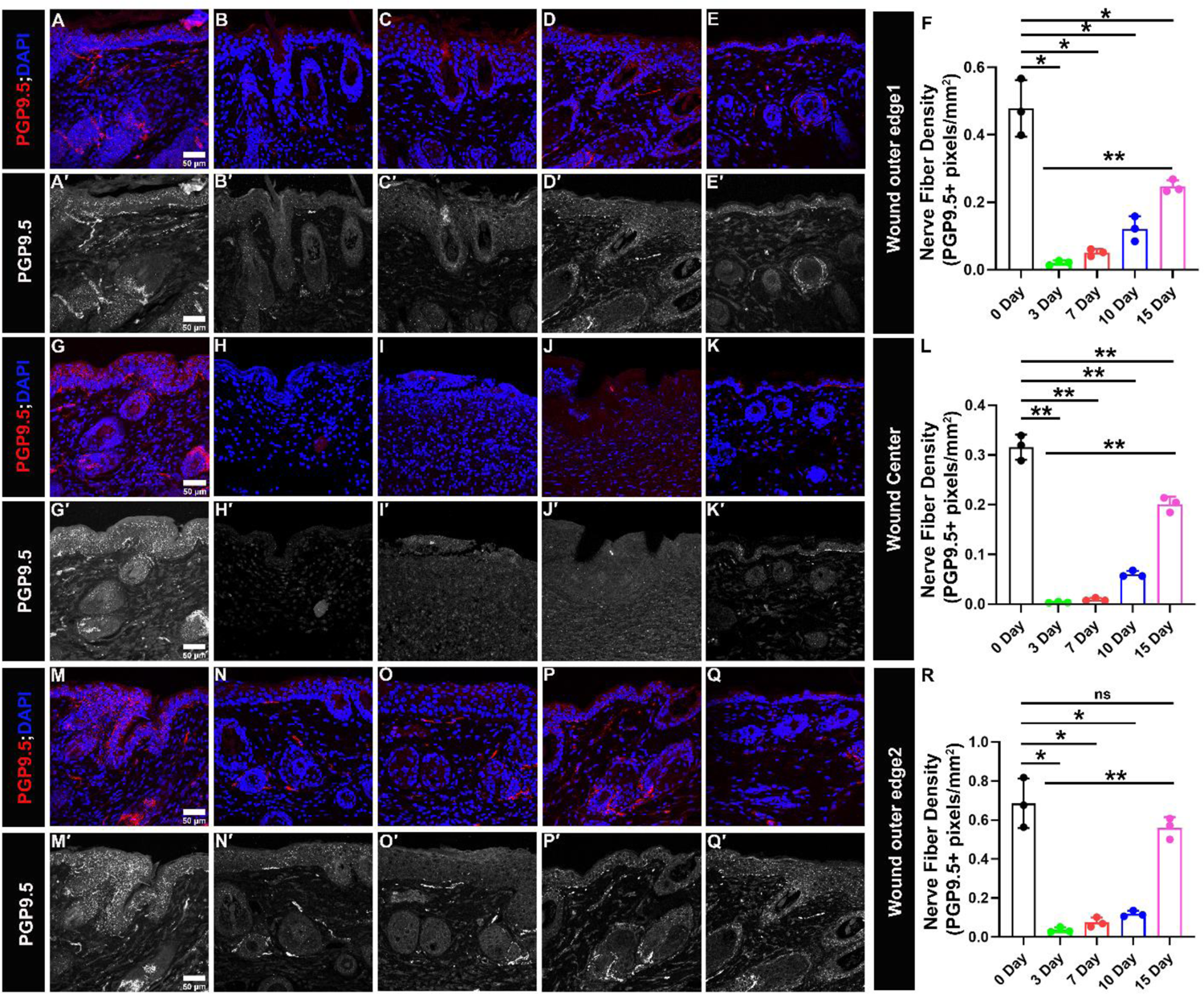
Gradual increase in Re-innervation at lateral wound edges and wound center. PGP9.5 is a pan-neuronal marker (in red) and DAPI stains nuclei of cell (in blue). (A′-E′, G′-K′, M′-Q′) Split images (in grey) show PGP9.5 immunoreactivity. (A-E & A′-E′) Immunoreactivity to PGP9.5 at wound edge 1 for skin samples (A, A′) Uninjured (B, B′) day 3 (C, C′) day 7 (D, D′) day 10 (E, E′) day 15 (F) Quantification of skin innervation at wound edge 1. (G-K & G′-K′) Immunoreactivity to PGP9.5 at wound center for skin samples (G, G′) Uninjured (H, H′) day 3 (I, I′) day 7 (J, J′) day 10 (K, K′) day 15 (L) Quantification of skin innervation at wound center. (M-Q & M′-Q′) Immunoreactivity to PGP9.5 at wound outer edge 2 for skin samples (M, M′) Uninjured (N, N′) day 3 (O, O′) day 7 (P, P′) day 10 (Q, Q′) day 15 (R) Quantification of skin innervation at wound edge 2. All quantification data are represented as mean ± SD, n = 3 wounds from three mice in each group, *P < .05, **P < .001, ns is non-significant. Scale bar = 50 μm.

### 2.5. Reepithelization

After fixation, the wound tissue was paraffin-embedded, sectioned to 5 μm and H&E stained for determination of wound re-epithelialization as described previously^29, 35^. Briefly, BioRevo BZ-9000 inverted microscope (Keyence, Osaka, Japan) was used to image all the histological sections. Measurements were done by an investigator blinded to experiment with the BZ-II viewer and analyzer (Keyence, Japan). The absence of underlying adipose tissue and hair follicles defines wound edges and wound healing is determined by the re-epithelialization of the epidermis layer^36–37^. The outgrowth of the newly formed epidermis was tracked manually from the wound edges and the percentage of the combined length of the re-epithelialization to the total length of the wounds was calculated.

### 2.6. Statistics

Statistical analysis was performed using an unpaired, two-tailed student *t*-test as described previously^38–39^. Data are expressed as mean ± SD. A *P*-value less than 0.05 was considered statistically significant.

## 3. Results

### 3.1. PGP9.5 as a specific neuronal marker

Anti-PGP9.5 antibody labels UCHL1/PGP 9.5 protein in the tissue sections (Figure 3,4). The protein is highly conserved and localized in neurons and is considered as a pan-neuronal marker that stains both sensory as well as autonomic nerves ^40–41^. To deduct background signal, we carried out experiments with negative control for every sample omitting the primary antibody (Supplementary figure 1). The background signals from the negative control are determined and used for generating a cutoff window for quantification of the true PGP9.5+ signals during MATLAB-based quantification.

To verify the authenticity of nerve fiber staining with PGP9.5, we also stained skin samples with another neuronal marker, β-III Tubulin that colocalized with the PGP9.5 staining (Supplementary figure 2). PGP9.5 staining is used widely and considered the gold standard for labeling skin innervation^40, 42^, so throughout the study, we quantified PGP9.5 immunofluorescence staining for the detection of nerve fibers.

### 3.2. Automated image processing and quantification of skin innervation

The quantification method used in this study employed an automated Matlab-assisted image analysis tool with a deep neural network to pre-process images and determine the range of skin innervation during different stages of wound healing. The use of the deep neural network ensured that noisy pixels did not affect the calculation of total neuronal coverage (Figure 2). Next, it was critical to determine the threshold value for identifying neurons because, if a threshold is too high, some neurons might not be detected, resulting in an underestimation of the neuronal coverage. Conversely, if the threshold is too low, non-neuronal elements in the image may be mistakenly identified as neurons, leading to an overestimation of neuronal coverage. To address this issue, we employed a statistical approach to determine the appropriate threshold value. First, we saved the R values of every pixel in the image to a list. We then calculated several statistical measures, including the minimum, maximum, mean, median, and interquartile range. Outliers were identified based on this statistical analysis and excluded from further consideration. To determine whether a pixel contained a neuron or not, we used the distance of the R-value from the third interquartile (Q3). Pixels with a closer distance to the maximum value in comparison to Q3 were selected as neurons. This method ensured that the threshold value was based on a statistically robust approach, which increased the accuracy of the quantification method. Furthermore, this approach required minimal manual intervention, making it suitable for large-scale studies and a reliable method for quantifying skin innervation during wound healing.

### 3.3. Wounding reduced nerve fiber density

Immunoreactivity was detected in intraepidermal and dermal nerve fibers and cells. The positively stained nerve fibers are quantified for the whole wound (Figure 3; Table 1) and separately for lateral wound boundaries, and wound center (Figure 4; Table 1). The nerve fiber density individually for the epidermis and dermis is also quantified respectively (Supplementary figure 3; Table 1). The total nerve fiber density for uninjured skin (unwounded) is found to be 0.92 ± 0.23, out of which 0.64 ± 0.21 are found in the epidermis and 0.27 ± 0.07 in the dermis. Comparaed to uninjured skin, as expected after creating a wound there is a considerable reduction in nerve fiber density throughout the wound bed (WB) (Figure 3; Table 1).

**Table 1.** Nerve fiber density at wound bed, wound outer edge 1, wound center, and wound outer edge 2 on day 0, 3, 7, 10 and 15 of healing.

| **Wound Bed** |  |  |  |  |  |  |  |  |  |  |  |  |  |  |
| --- | --- | --- | --- | --- | --- | --- | --- | --- | --- | --- | --- | --- | --- | --- |
| **Epidermis** |  |  |  |  | **Dermis** |  |  |  |  | **Whole wound** |  |  |  |  |
| **0 Day** | **3 Day** | **7 Day** | **10 Day** | **15 Day** | **0 Day** | **3 Day** | **7 Day** | **10 Day** | **15 Day** | **0 Day** | **3 Day** | **7 Day** | **10 Day** | **15 Day** |
| 0.64<br>±<br>0.21 | 0.0±<br>0.0 | 0.0±<br>0.0 | 0.08<br>±<br>0.00<br>4 | 0.44<br>±<br>0.04 | 0.27<br>±<br>0.07 | 0.02±<br>0.01 | 0.04±<br>0.03 | 0.10<br>±<br>0.00<br>4 | 0.16<br>±<br>0.00<br>7 | 0.92<br>±<br>0.23 | 0.02<br>±<br>0.01 | 0.04<br>5 ±<br>0.00<br>3 | 0.18<br>±<br>0.01 | 0.6±<br>0.05 |
| **Wound outer edge 1** |  |  |  |  |  |  |  |  |  |  |  |  |  |  |
| **Epidermis** |  |  |  |  | **Dermis** |  |  |  |  | **Whole wound** |  |  |  |  |
| **0 Day** | **3 Day** | **7 Day** | **10 Day** | **15 Day** | **0 Day** | **3 Day** | **7 Day** | **10 Day** | **15 Day** | **0 Day** | **3 Day** | **7 Day** | **10 Day** | **15 Day** |
| 0.4±<br>0.1 | 0.01<br>±<br>0.01 | 0.04<br>±<br>0.01 | 0.1±<br>0.04 | 0.2±<br>0.02 | 0.08<br>±<br>0.02 | 0.01±<br>0.001 | 0.01±<br>0.001 | 0.01<br>±<br>0.01 | 0.05<br>±<br>0.00<br>3 | 0.48<br>±<br>0.08 | 0.02<br>±<br>0.01 | 0.05<br>±<br>0.01 | 0.12<br>±<br>0.04 | 0.25<br>±<br>0.02 |
| **Wound center** |  |  |  |  |  |  |  |  |  |  |  |  |  |  |
| **Epidermis** |  |  |  |  | **Dermis** |  |  |  |  | **Whole wound** |  |  |  |  |
| **0 Day** | **3 Day** | **7 Day** | **10 Day** | **15 Day** | **0 Day** | **3 Day** | **7 Day** | **10 Day** | **15 Day** | **0 Day** | **3 Day** | **7 Day** | **10 Day** | **15 Day** |
| 0.2±<br>0.02 | 0± 0 | 0± 0 | 0±0 | 0.09<br>±<br>0.02 | 0.13<br>±<br>0.02 | 0.01±<br>0.001 | 0.009<br>±<br>0.003 | 0.06<br>±<br>0.01 | 0.11<br>±<br>0.01 | 0.31<br>±<br>0.03 | 0.00<br>3±<br>0.00<br>1 | 0.00<br>8±<br>0.00<br>3 | 0.06<br>±<br>0.00<br>6 | 0.2±<br>0.01<br>5 |
| **Wound outer edge 2** |  |  |  |  |  |  |  |  |  |  |  |  |  |  |
| **Epidermis** |  |  |  |  | **Dermis** |  |  |  |  | **Whole wound** |  |  |  |  |
| **0 Day** | **3 Day** | **7 Day** | **10 Day** | **15 Day** | **0 Day** | **3 Day** | **7 Day** | **10 Day** | **15 Day** | **0 Day** | **3 Day** | **7 Day** | **10 Day** | **15 Day** |
| 0.28<br>±<br>0.07 | 0.02<br>±<br>0.01 | 0.04<br>±<br>0.00<br>3 | 0.10<br>±<br>0.00<br>4 | 0.16<br>±<br>0.00<br>9 | 0.13<br>±<br>0.01 | 0.006<br>±<br>0.002 | 0.009<br>±<br>0.004 | 0.04<br>±<br>0.01 | 0.10<br>±<br>0.00<br>2 | 0.67<br>±<br>0.13 | 0.03<br>6±<br>0.01<br>3 | 0.07<br>4±<br>0.02<br>4 | 0.11<br>9±<br>0.01<br>5 | 0.56<br>±<br>0.05 |

### 3.4. Innervation gradually appears and reaches a level close to intact skin-15 days after wounding

On day 3 of wound healing, nerve fiber density in the wound bed is found to be 0.02 ± 0.01 which is significantly less than the unwounded skin (0.02 ± 0.01 Vs 0.92 ± 0.23). The nerve fiber density reported is found only in the dermis (0.02 ± 0.01), yet the epidermis has not regenerated in the wound bed, so the intraepidermal nerve fiber density value is 0. On day 7, the nerve fiber density increases slightly to 0.045 ± 0.003. The intraepidermal nerve fiber density is still 0. The wound sections on day 10 show some traces of intraepidermal nerve fibers, the nerve fiber density quantified is 0.080 ± 0.004. The nerve fiber density in the dermis, 0.10 ± 0.004, also steadily shows an increasing trend. Both intraepidermal and dermis nerve fiber density counts together to 0.18 ± 0.01 on day 10, which is still significantly less compared to unwounded skin (0.18 ± 0.01 Vs 0.92 ± 0.23). Interestingly, on day 15 of healing there is a huge increase in intraepidermal nerve fiber density, 0.44 ± 0.04. The nerve fiber density in the dermis also substantially increases to 0.16 ± 0.007. On day 15, the changes in nerve fiber density for the whole wound bed compared to unwounded skin (0.6 ± 0.05 Vs 0.92 ± 0.23) are non-significant (Figure 3 and Supplementary figure 3) and interestingly significant when compared to day 3 of healing (0.6 ± 0.05 Vs 0.02 ± 0.01). The data narrates that there is a gradual increase in nerve fiber density through the time series of wound healing, and there is a significant re-innervation and values start reaching close to normal from day 15 of healing onwards.

### 3.5. The wound center and wound edge show a similar trend in re-innervation

We also quantified nerve fibers separately for wound edges and wound center on days 3, 7, 10, and 15 of wound healing and compared it with the innervation of unwounded skin (Figure 4 and Supplementary figure 3; Table 1). For unwounded skin the values obtained at outer edge 1 are: intraepidermal (IE) = 0.4 ± 0.1; dermis (D)= 0.08 ± 0.02; both together (IE+D) =0.48 ± 0.08. It is evident that on day 3, 7 and 10 there is a significant decrease in the IE and D nerve fiber density compared to uninjured skin (Table 1). As expected, the nerve fiber density steeply increased on day 15 of healing: IE= 0.2 ± 0.02; D= 0.05 ± 0.003; IE+D= 0.25 ± 0.02. The values show non-significant change compared to uninjured skin except for both (IE+D) where nerve fiber density at outer edge 1 is still significantly lesser compared to uninjured skin (0.25 ± 0.02 Vs 0.48 ± 0.08). Also, the change found is significant compared to 3 days for IE, D and both taken together (IE+D) (Figure 4 and Supplementary figure 3; Table 1). Therefore, data signify that at outer edge 1 of the wound considerable level of re-innervation takes place up to day 15 of healing, however, total nerve fiber density cannot reach to the normal level i.e. the values are significantly less compared to the uninjured skin. Contrary to which at outer edge 2 significant reinnervation happens but total nerve fiber density at day 15 also show non-significant change compared to uninjured skin at IE, D, and both together (IE+D) (Figure 4 and Supplementary figure 3; Table 1). Whereas, wound center behaves like outer edge 1 where a significant re-innervation happens by day 15 but the nerve fiber density compared to uninjured skin (0.2 ± 0.015 Vs 0.31 ± 0.03) show significantly lower values. Stating that a considerable increase in nerve fibers at the wound outer edge 1 and wound center is still to be expected after day 15.

### 3.6. Re-innervation of the wound correlates strongly with re-epithelialization

Denervation has a detrimental effect on cutaneous wound healing. Severing the nerves hinders cutaneous wound healing, and sympathetic denervation of the skin delays re-epithelization^43^. Re-epithelization is defined by the epithelial cells migrating and growing over the wound bed and complete epithelial covering of the wound is a criterion to evaluate if a wound has healed properly ^29, 44^. We investigated the correlation between nerve fiber density and re-epithelization on days 3,7,10, and 15 of healing. Our data demonstrate a strong positive correlation (R^2^=0.933) between the two (Figure 5). This substantially corroborates our hypothesis that the regeneration of nerve fibers is critical for proper wound healing in time.

**Figure 5:**
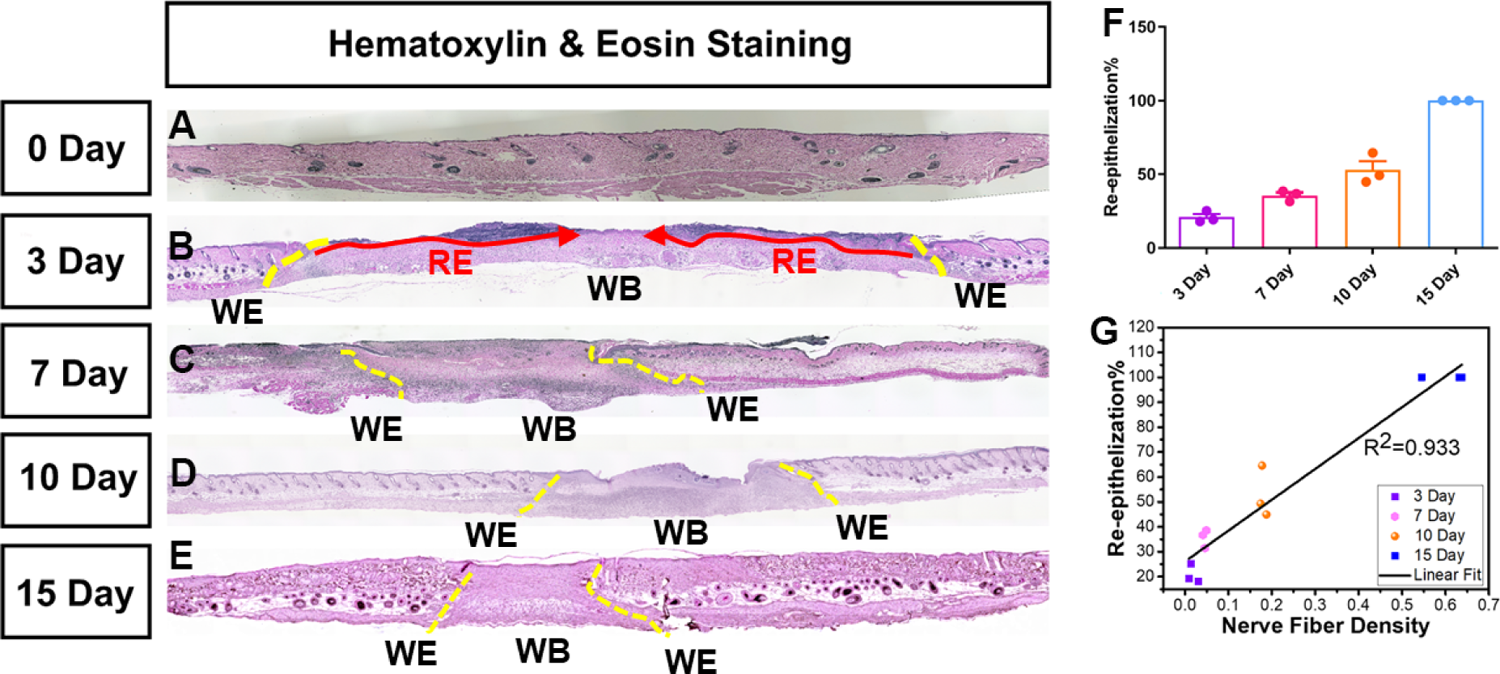
Positive correlation between re-innervation and re-epithelization. A representative image of H&E staining of the skin sample collected on (A) day 0 (B) day 3 (C) day 7 (D) day 10 (E) day 15. The original wound edge (yellow dashed lines) on each side is determined by the absence of subdermal adipose tissue. Re-epithelialization (red arrows) is defined by epithelial cell growth above the Integra matrix. (F) quantification of re-epithelization (G) correlation between nerve fiber density and re-epithelization at day 3,7,10 and 15 of wound healing. R^2^=0.933 show strong positive relation. Here abbreviations represent WE: wound edge; WB: wound bed; RE: Re-epithelization.

## 4. Discussion

The skin is densely innervated with complex architecture and network of cutaneous nerves, which are present in both epidermis and dermis. The degree of innervation has a direct effect on all the overlapping stages of wound healing^1^, and previously it has been reported that denervated wounds take longer time to heal^45^. Therefore, precise quantification of nerve fiber density during different stages of wound healing becomes critical. Previously PGP9.5 staining is considered as a gold standard for the quantification of nerves in skin samples in mammals^40, 42^. However, precise quantification of cutaneous nerves is challenging because of background noise and a different pixel intensity of PGP9.5+ neurons throughout epidermis and dermis. Therefore, to solve the problem we used powerful deep neural network, DnCNN, for pre-processing (de-noising) of the IHC-images, followed by stringent statistical methods to formulate a threshold boundary, which is broad enough to include all the features of interest and strict enough to exclude background, so that PGP9.5+ pixels are precisely quantified in cutaneous wounds. Applying our newly developed technique, automated Matlab-assisted tool aided with DnCNN, we quantified skin innervation in Female C57BL/6 mice during normal wound healing at days 3,7,10 and 15. The data show that (1) the skin wound causes substantial reduction in nerve fiber density, which significantly increased by day 15 of wound healing (2) One of the wound edge and wound center still have significantly less nerve fiber density compared to uninjured skin at day 15 of wound healing (3) re-epithelization and innervation share a strong positive correlation (R^2^=0.933).

We found that on day 3 and day 7 nerve fiber density is almost negligible when compared to the unwounded skin (Figure 3; Table 1). From day 10 onwards the nerve fiber density starts increasing marginally but reaches a significant level at the later stage of wound healing on day 15. At day 15, comparing the nerve fiber density for the whole wound or individually for the epidermis as well as dermis shows non-significant change when compared to the uninjured skin for the respective skin regions. Moreover, interestingly the values are significantly higher when compared to the initial stages of wound healing, day 3, at respective skin regions (Figure 3; Supplementary figure 3; Table 1). This trend together suggests that re-innervation of the wound is initiated at the later stages of wound healing and starts becoming substantial from day 15 onwards during normal wound healing conditions in mammals. The data seems logical because for re-innervation, proliferation of neuronal cells needs to be initiated, followed by orchestrated phenomena to develop new cutaneous nerves^46–50^, and the proliferation stage of neuronal cells might overlap with the proliferation stage of wound healing, which begins approximately at day 3 and lasts for a couple of weeks. Thus, re-innervation also starts appearing substantially from day 10 onwards and shows a significant increase by day 15. Quantifying nerve fiber density separately for wound edges and wound center showed an interesting trend. At outer edge 2, similar to the whole wound, the total nerve fiber density at day 15 showed non-significant change compared to uninjured skin at IE, D, and both together (IE+D) (Figure 4 and Supplementary figure 3; Table 1). However, for the outer edge 1 and wound center total nerve fiber density cannot reach to the pre-injury level i.e., the values are still significantly less compared to uninjured skin for the whole wound bed (Figure 4). Suggesting that nerve fibers are continuously innervating the cutaneous wound even beyond day 15 of wound healing. Also, on day 15 of healing the nerve fiber density at outer edge1 is less compared to outer edge2 (0.25 ± 0.02 Vs 0.56 ± 0.05) (Figure 4). This could be due to the difference in distance between the two outer edges of the skin wound from the cell bodies located in the dorsal root ganglia from where the cutaneous sensory nerves originate^51^. Finally, we have been able to find a strong positive correlation (R^2^=0.933) between re-epithelization and nerve fiber density during time-series of wound healing, which not only corroborates the fact that the regeneration of nerve fibers is critical for proper wound healing in time but also validated our technique of using automated Matlab-assisted tool aided with DnCNN for denoising to precisely capture PGP9.5 + pixels, and thus calculate nerve fiber density.

Overall, although our method does not elucidate the morphological characteristics of cutaneous nerves in three dimensions, it does, simply and economically, address the differences in the extent of skin innervation at different stages of wound healing. The statistical approach presented in this study offers a useful alternative to previously described methods ^20–26^ and machine learning-based techniques for quantifying skin innervation during wound healing. By using this approach, researchers can obtain accurate and reliable results on high throughput experiments, while reducing the amount of time, labor, and resources needed to obtain reliable outcome measurements.

## 5. Conclusion and Future Directions

This study reports utilizing automated deep-learning tools to reliably quantify skin innervation in the wound bed as well as at the wound center and wound edges in mammals. The sensitivity of the technique to de-noise IHC images made it possible to exclusively quantify intraepidermal and dermis nerve fiber density for respective wound areas (bed, center, and lateral edges). This data-driven approach generates critical information that can provide a predictive model for applications in precision medicine in wound healing. Furthermore, combining a statistical approach with deep learning eliminated the need to manually label data as is needed in traditional machine learning approaches to generate training data. The denoise method used in this study had already been trained on biological images, eliminating the need for further training on wound images. This means that researchers can save time and resources that would otherwise be spent on data collection and annotation. The data generated shows that there is a gradual increase in nerve fiber density throughout the wound bed as well as at the wound edges, with maximum value reaching on day 15 that indicates a significant trend in re-innervation when compared to day 3 of wound healing. Additionally, on day 15 of healing nerve fiber density for whole wound bed, and at wound edge 2 reaches close to the uninjured skin. However, interestingly at wound outer edge 1 and wound center the total nerve fiber density at day 15 of healing is still significantly less compared to the uninjured skin asserting that re-innervation is still a continuous process beyond day 15 of healing and is important for the complete healing of the wound. A positive correlation between the increase in nerve fiber density and re-epithelization further supports the importance of cutaneous nerve fibers in wound healing.

Many treatments have been reported to speed up and/or improve the quality of wound healing. Electric stimulation, for example, has shown promising results in promoting innervation in human cutaneous wounds^52–54^. Many bandages e.g., Procellera claim to deliver electric fields to promote wound healing^54^. Our approach and methodology developed in this paper can assist the quantitative determination of nerve fiber density in space through the time of wound healing and facilitate assessing the effects on wound treatment. This high throughput method can be adopted for the quantification of innervation in various skin pathologies in addition to wound healing, and for the quantification of innervation of other tissues and organs.

## Acknowledgements

We will like to thank Prof. Qizhi Gong for training us with the Confocol microscopy.

## 7. Author Contributions

A.S.M, R.R.I, M.Z, M.G, designed the study. A.S.M, S.T, C.R, D.F, A.G, H.Y, performed the experiments. R.R.I, M.Z., M.G., contributed the resources. A.S.M, S.T, R.R.I, M.Z., M.G., analyzed the data. A.S.M, M.Z, wrote the manuscript with input from all authors. All authors read and approved the final manuscript.

## 9. Data availability

The authors confirm that the data supporting the findings of this study are available within the article or its supplementary materials.

---

**Supplementary Figure 1:**
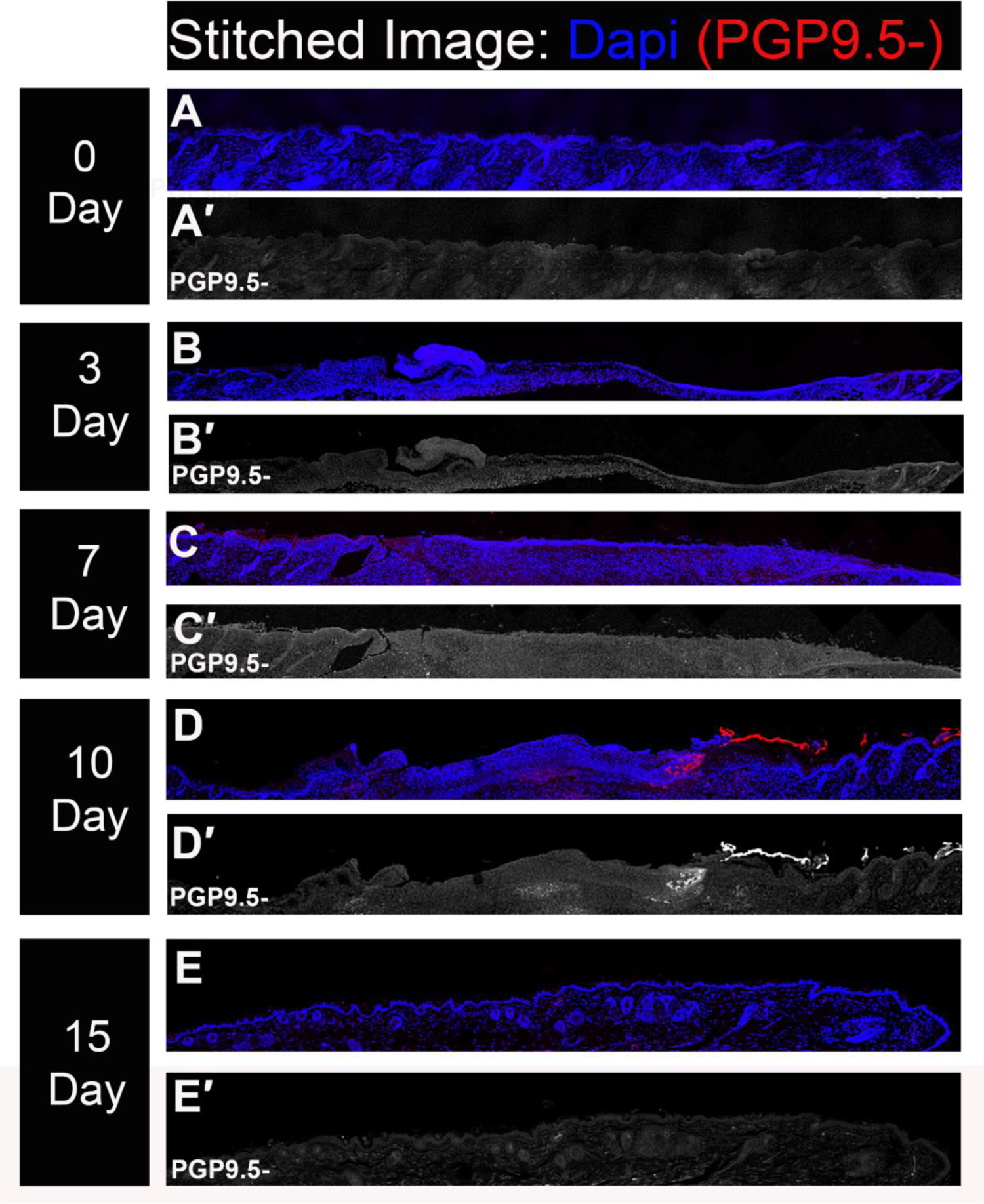
Stitched Immunohistochemistry images of 5 μm vertical sections of punch biopsies as a negative control for PGP9.5. PGP9.5 is a pan-neuronal marker and DAPI stains the nuclei (In blue). (A) Uninjured skin. Skin samples were collected on (B) day 3 (C) day 7 (D) day 10 (E) day 15. The red color is autofluorescence from a dead skin flap.

**Supplementary Figure 2:**
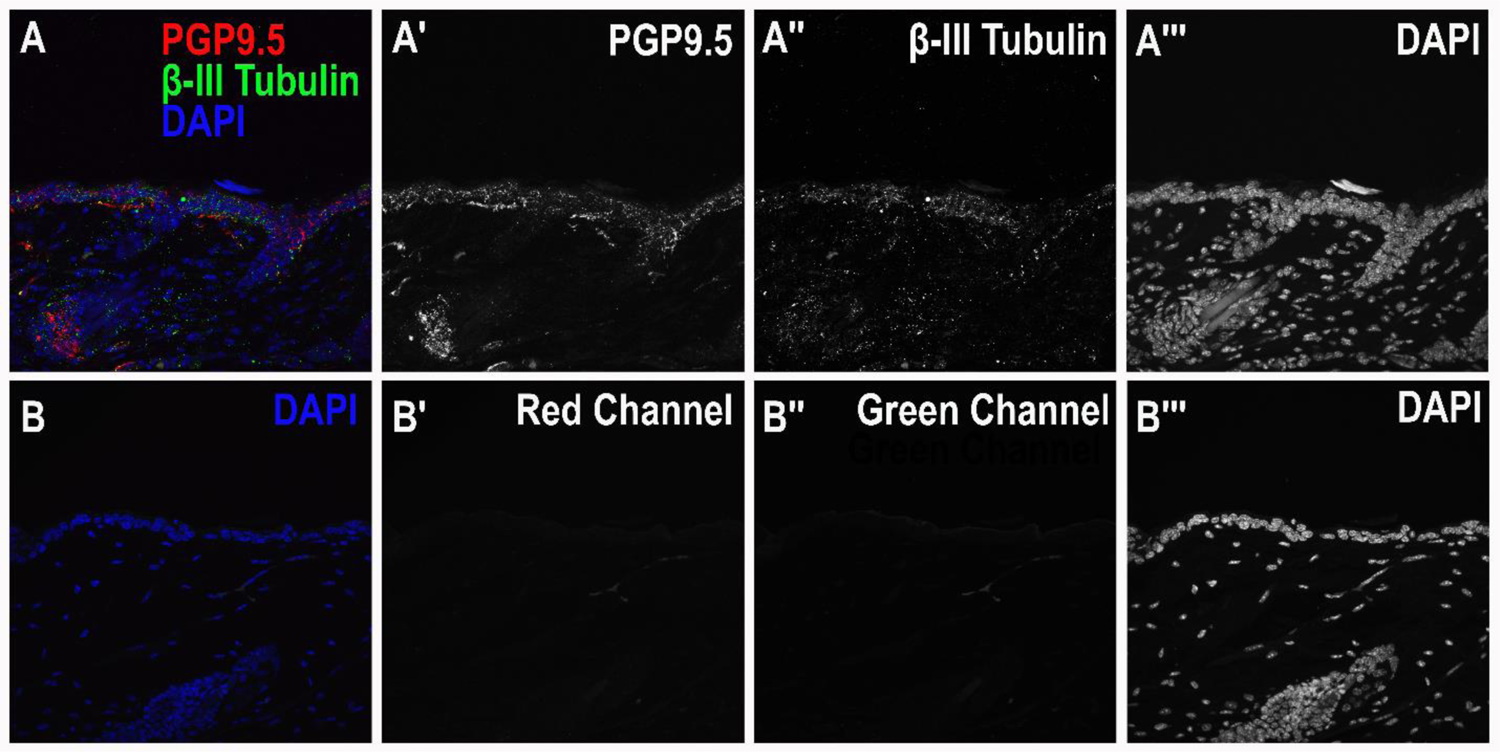
Immunoreactivity to PGP9.5 and Beta-III tubulin as double staining for neurons. (A, B) A representative example of uninjured skin showing double staining with PGP9.5 and β-III tubulin (A,A′,A′′,A′′′) Positive staining (B,B′,B′′,B′′′) Negative control.

**Supplementary Figure 3:**
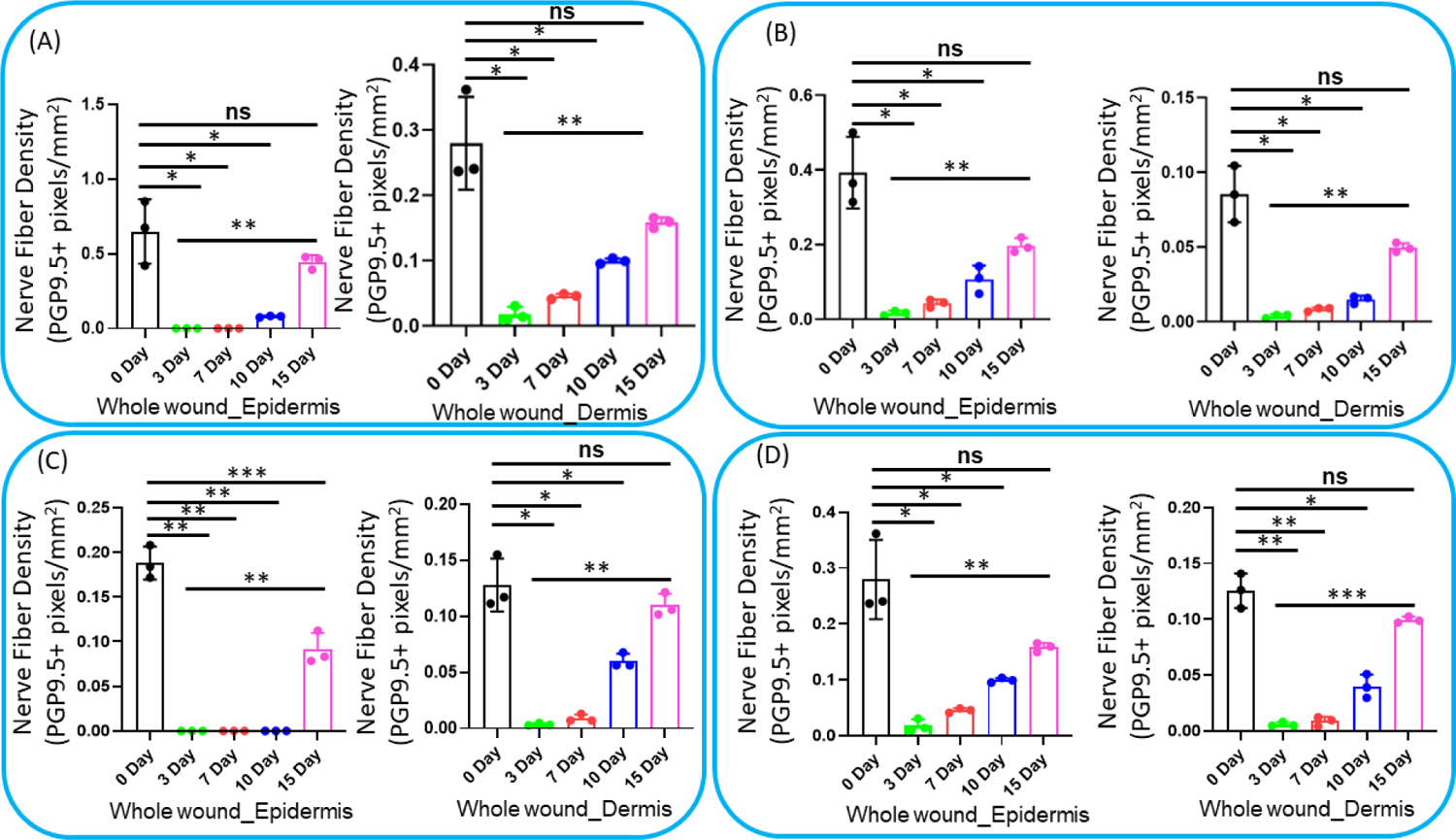
Intraepidermal and dermis innervation. Quantification of innervation in epidermis and dermis for (A) whole wound (B) Wound outer edge 1 (C) Wound center (D) Wound outer edge 2. All quantification data are represented as mean ± SD, n = 3 wounds from three mice in each group, *P < .05, **P < .001, ***P < .0001, ns is non-significant.

